# Quantitative horizon scanning identifies high-priority wood-boring beetle pests for regulatory review in Canada

**DOI:** 10.64898/2026.01.05.697691

**Authors:** JR Stinziano, W Zheng, M Abergel, M Damus, A Ameen, H Cumming, R Dimitrova

## Abstract

Invasive wood-boring beetles are a significant threat to forests, imposing high economic and environmental costs. With their high global biodiversity, it is difficult to assess all wood-boring beetles’ potential to invade a particular region. Horizon scanning can create short-lists of high priority pests for risk assessment by National Plant Protection Organizations; however, it is often resource-intensive, creating potential blind spots due to resource constraints. Here, we use a multi-layered horizon scan, including self-organizing maps and climate suitability modelling combined with rapid risk assessment, to produce a short-list of wood-boring beetles for regulatory review in Canada. This method relies on open-source data and could be applicable to any country. From an initial list of 10,824 species with available georeferenced observations, our method yielded a short-list of 24 species. The method led to the regulation of two additional wood-boring beetles in Canada, demonstrating its real-world applicability in accelerating the regulation of quarantine pests. Several already-regulated species were also identified, suggesting consistency with existing risk identification and assessment procedures. The limitations on the method are the availability of biologically meaningful species occurrence data, relevance of climate norms for species’ establishment, and availability of data for rapid risk assessment.

## Introduction

Global trade is a primary introduction pathway of invasive alien species (IAS) that may have previously been bound by natural borders (Westphal, 2008). The economic cost of IAS is estimated at >US $1.26 trillion in North America over the last 50 years (Crystal-Ornelas et al., 2021). Costs of managing IAS are proportional to time since introduction. Prevention has the lowest costs, and mitigation and management costs increase as IAS spread until it is no longer economical to stop or reduce spread (Cuthbert et al., 2022). Therefore, early identification of potential invaders before establishment is important for preventing their introduction.

Early threat identification is difficult for many reasons, including but not limited to lack of data on species, changes in pest behaviours, or niche expansion into new environments (Ricciardi et al., 2017; Antoniou et al., 2024). Within National Plant Protection Organizations (NPPOs) responsible for regulating and controlling IAS, formal risk assessment procedures identify whether a species is a potential pest threat (IPPC, 2021a; IPPC, 2021b; IPPC, 2021c). These processes look at likelihood of entry based on known pathways, likelihood of establishment based on environmental suitability and host availability, ability to spread, and potential environmental and economic costs associated with the species establishing in a particular country. These formal risk assessments are comprehensive, thorough, and time-consuming, meaning only a limited number of species can be assessed in any given year. Techniques to prioritize potential pests may improve efficiency of the regulatory continuum and enhance plant protection efforts worldwide.

Horizon scanning is used to identify potential threats or impactful changes that could happen in the near term (EEA, 2023). Typical techniques involve review of the literature, expert panels, and relatively qualitative assessment of species to determine whether they represent a threat to plant health, with the desired result being a short-list of species to carry forward to formal risk assessment (Kendig et al., 2022; Antoniou et al., 2024). These techniques are labour-intensive and suited to jurisdictions with sufficient resources to conduct the entire workflow of activities. However, given trade globalization (Ricciardi et al., 2017) and increasing uptake of citizen science initiatives for reporting pests (Howard et al., 2022), these activities could be augmented with reproducible and (semi-) quantitative algorithms.

Here we present a method for quantitative horizon scanning to identify potential pests for formal risk assessment. Our objective was to develop methods that would produce a short-list using open-source data and modelling techniques, to offload effort in horizon scanning from a human operator to computers. Our case study focuses on wood-boring beetle threats to Canada, as wood-boring beetles have relatively well-defined taxa, allowing us to test methodology on a subset of available data, and they represent serious economic and environmental threats to Canada’s forestry industry and ecology. Even with international systems in place to reduce movement of wood-boring beetles in trade (ISPM 15: FAO 2018), interceptions of these species are occurring with increasing frequency due to increasing trade (Haack et al. 2022; Zahid et al. 2008).

## Materials & Methods

### Species occurrence data

We targeted wood-boring beetles (Order: Coleoptera). Nine families of true wood-boring beetles have been identified (USDA, 2018) viz., Anobiidae, Bostrichidae, Brentidae, Buprestidae, Cerambycidae, Curculionidae, Lymexylidae, Oedemeridae, and Zopheridae. Curculionidae is very diverse, with wood-boring beetles found within 37 genera in the subfamily Scolytinae or 246 genera in the subfamily Platypodinae (Hulcr et al., 2015; Wood, 1993).

We obtained occurrence data for species in the above groups from GBIF (GBIF, 2024) using {rgbif} (Chamberlain et al., 2024; Chamberlain and Boettiger, 2017), resulting in 10,824 species. We further refined this list by excluding species with observations in Canada (Bousquet et al., 2013), which reduced the list to 10,028 species.

### Trade data

Global trade has introduced serious damaging invasive species into Canada such as the emerald ash borer (NRCan, 2024) and the Asian longhorned beetle (ISC, 2024). Wood-boring beetles could enter Canada on any number of commodities if commodities are accompanied by wooden packaging materials. Trade data were obtained from Global Trade Tracker (Global Trade Tracker, 2023). We compiled a list of 164 countries that have exported any commodity, of any quantity, to Canada between 2018 and the date the data was pulled (July 2, 2023).

Total trade volume was calculated as mean annual sum of all imports to Canada from 2018 to 2023. For each species, a trade score (TS) was calculated as:

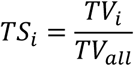

Where TS_i_ is the Trade Score for species i and is bounded by 0 and 1, TV_i_ = trade volume for Canadian trading partners in which species i is found, and TV_all_ is trade volume for all trading partners.

Combining the trade data with the origin country from the occurrence data, a weighted country score was calculated as:

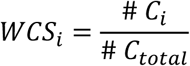

Where WCS_i_ is the weighted country score for species i, # C_i_ is the number of countries in which species i is known to occur, and # C_total_ is total number of countries.

### Climate data

Climate data were obtained from CHELSA (Climatologies at high resolution for the earth’s land surface areas; Kerger et al., 2017) for 1981-2010 (Table 1). All climate data was aggregated to 10 arc-minutes, using = *aggregate* from {terra} (Hijmans, 2024).

**Table 1.**
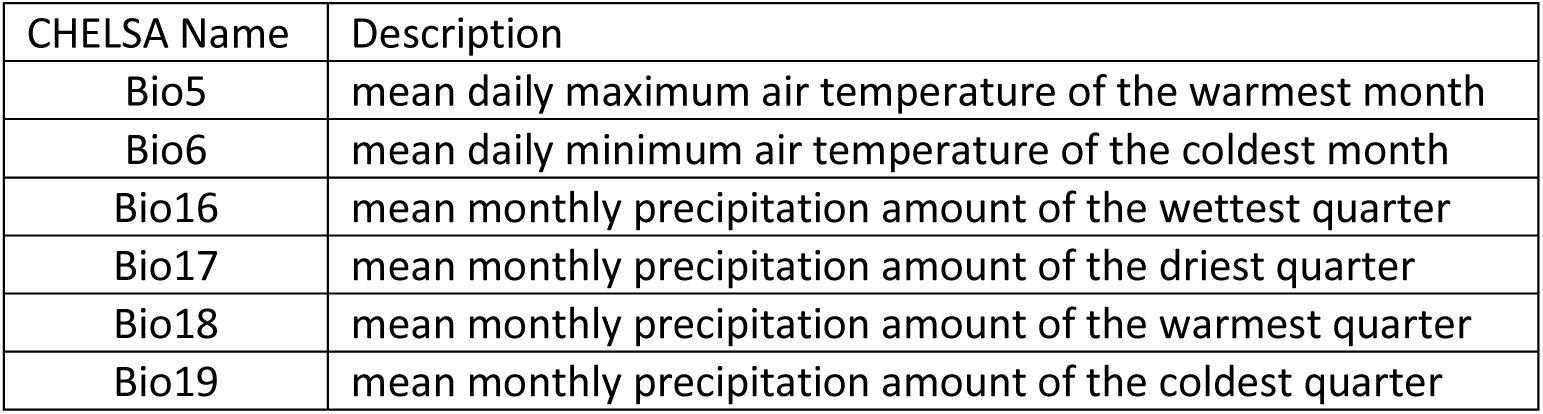
Climate variables used for climate matching. See Table S1 for all variables considered.

We ran a correlation analysis to select climate variables for climate matching. Correlation matrices were created using *corrplot* from {corrplot} (Wei and Simko, 2021). Highly correlated variables can influence the covariance structure and make interpretation difficult (Venette, 2015), therefore, a correlation matrix of 14 variables was examined. One variable from each correlated pair was stepwise removed, and correlation analysis re-run, until all remaining variable correlation values were <0.8 (Figures S1 and S2). We looked at thresholds of 0.7 and 0.8, using a threshold of 0.8, rather than 0.7 as recommended by Dormann et al. (2013), because a threshold of below 0.8 removed some variables considered *a priori* as variables of biological significance (bio5 and bio6).

### Climate matching

Climate matching identifies climate niches potentially suitable for species based on similarity to climates found in the species’ current range. The Climatch algorithm, shown below, allows for these climate comparisons (Crombie et al. 2008).

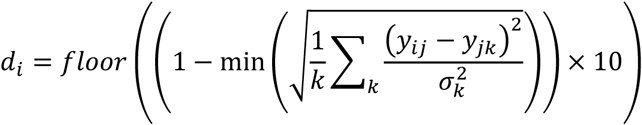

Where d is Climatch score, i and j represent two sites over which the calculation is made, k is the k^th^ climate variable, y is the climate variable value, and σ^2^ is the climate variable standard deviation.

Climatch has been validated as a consistent predictor of invasion risk across a range of taxa (e.g. Bomford 2008; Bomford et al. 2009). We used {climatchR} for calculating Climatch scores (Erickson et al., 2022). Briefly, each cell within a raster of Canada is assigned a score between 0 – 10, where 10 is a perfect climate match between source locations and the target point and 0 is no match. The number of cells with a score of ≥6 is divided by the total number of cells to calculate climate suitability score. This cut-off was determined by looking at minimum Climatch scores required to correctly predict 98% of observations of wood-boring beetles already present in Canada (Stinziano, unpublished).

### Pest status, hosts, and forest impact data

Pest status was determined based on whether a species was regulated by one of Canada’s largest trading partners by trade volume, namely the US (IPPC, 2024), China (IPPC, 2024), Mexico (IPPC, 2024), Japan (IPPC, 2024), and Europe (EPPO, 2024). Pest Score, PS, was set to 1 if the species was regulated as a pest.

Pest-host relationship data were obtained from EPPO (EPPO, 2024). The genera of potential hosts for species with climate or SOM suitability were cross-referenced with major tree genera in Canada.

Host Score (HS) was calculated as:

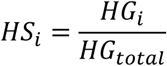

Where HS_i_ is host score for species i and is bounded by 0 and 1, HG_i_ is number of host genera present, and HG_total_ is total number of tree genera present in Canada.

To calculate potential forest impact, merchantable wood volume and % genus composition data for Canada’s forests were obtained from the NFI MODIS 250m resolution 2011 dataset (Beaudoin et al., 2017). Merchantable wood volume for each genus was estimated according to:

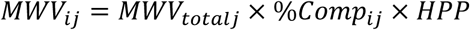

Where MWV_i_ is merchantable wood volume for genus i in m^3^ for pixel j, MWV_total_ is total merchantable wood volume in m^3^/ha for pixel j, %Comp is % composition of wood volume for genus i in pixel j, and HPP is a hectares per pixel conversion factor of 6.25 hectares per pixel.

Next, for each species, we extracted all values of MWV for pixels with a Climatch score exceeding 6 for each genus that was considered a host based on the EPPO database. A forest impact score was then calculated as:

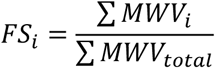

Where FS_i_ is forest impact score for species i and varies between 0 and 1, and ∑MWV_total_ is total merchantable wood volume for all of Canada. For species without known hosts, the FS was set to 0.

### Self-organizing maps

A self-organizing map (SOM) is an unsupervised artificial neural network (Kohonen, 1982) with an input layer and output layer (Roigé et al., 2016). Input neurons in this case are species assemblages of wood-boring beetles. SOMs capture similarity within regional species assemblages (input layer), positions them within a multidimensional space based on their similarity and projects them onto the output layer where species assemblages are clustered in a way that maximizes their similarity among geographic regions (Worner & Gevrey, 2006; Roigé et al., 2016). The output layer is a two-dimensional map comprising 5√n neurons arranged in a hexagonal lattice structure according to Vesanto’s rule for optimal SOM size, where n is the number of species (Vesanto et al., 2000). Each species is given a weight for each region, which ranges between 0 and 1 and can be interpreted as establishment risk values (Worner & Gevrey, 2006).

Species with weights closer to 1 are more closely associated with species assemblages of the target region, so a species with high weights that is absent from the target region could be interpreted as having potential to establish if introduced (Worner & Gevrey, 2006).

We ranked species for each Canadian province based on their SOM weight values, and kept species for evaluation that were absent in a province but present in other administrative areas located within the same neuron, as these species could be potential pests for that province (Worner & Gevrey, 2006; Roigé et al., 2016). Only species that occurred in at least two administrative regions, and with more than 2 observations (Worner & Gevrey 2006) were entered into the analysis. The final dataset was a matrix of 2650 administrative areas and 5529 species.

SOM analysis was performed using {Kohonen} (Version 3.0.12; Wehrens & Kruisselbrink, 2018, 2023). Data to assign state-level regions to occurrence points were from {rnaturalearth} (Massicotte & South, 2023). {aweSOM} was used for evaluating SOM quality for hyperparameter tuning with Kaski-Lagus error (Boelaert et al., 2022). Hyperparameter tuning led to a map size of 16 x16 (256 neurons) with hexagonal configuration and 500 iterations.

### SOM ensemble score

Since species risk values were estimated at the provincial and territorial level, we aggregated it to a national level. For each species in the SOM risk list:

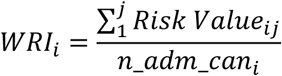

Where WRI_i_ is the weighted risk index for species i, and risk value is the SOM output for species i and administrative region j, and n_adm_can is the number of administrative areas in Canada where the species is identified as a potential pest for species i.

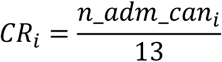

Where CR_i_ is the coverage ratio for species i, and 13 is the number of administrative areas in Canada. Two methods for calculating overall SOM score (SS) were:

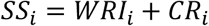

### Final scoring

Final scores were calculated using an additive and multiplicative approach (multiplicative calculation gave highly similar scores and is not shown). The additive calculation was:

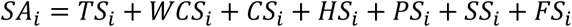

Where SA_i_ is the full additive score for species i.

### Short-list and validation procedure

To short-list species, we used segmented linear regression between score and rank to identify breakpoints in relationship with {segmented} (Muggeo, 2008) using the first 200 ranks, varying npsi from 1–3.

To validate the utility of results for NPPOs, we cross-referenced the species short-list with the CFIA regulated pest list (CFIA, 2024). A practical test to see if a horizon scan is effective at supporting risk assessment and pest regulation is to ask whether currently regulated pests are present in the short-lists and whether any additional species identified would be categorized as potential quarantine pests.

For short-listed species that were not already regulated, A. Ameen, H. Cumming, R. Dimitrova, and M. Damus applied CFIA’s pest categorization process to determine whether species meet the definition of a potential quarantine pest. We forwarded the list of potential quarantine pests to forestry program officers at the CFIA for regulatory review where regulatory decisions were made.

### Comparison of beetle families

We used a Kruskal-Wallis H test to determine whether there were differences in scores between beetle families, followed by Dunn’s test, using the Benjamini-Hochberg procedure to control for false discovery rate using {dunn.test} (Dinno, 2024).

### Analysis

All analyses were conducted using R (R Core Team, 2024).

## Results

Climatch identified 1,652 species with climate suitability in Canada, while SOM identified 489 species with establishment potential, for a total of 1,935 species (206 species overlapped). To determine whether overlap of species could be due to randomness, we randomly sampled input species lists for Climatch and SOM, taking 1,652 species from the Climatch input list and 489 species from the SOM input list and repeating the procedure 1,000 times. Mean overlap between the random lists was 116.7 ± 9.1 species. Assuming a normal distribution in number of overlapped species, the probability of getting 206 species overlapping randomly is < 1.3 X 10^−22^. Of 1,935 species, 824 are considered pests by at least one of EPPO, USA, China, Mexico, and Japan, and a total of 36 species are already regulated by Canada.

Score-rank relationships followed tailed distributions (Figure 1), with a rapid drop-off in scores with rank. Segmented regression produced a short-list of 24 species (Table 2). For the 10 species that were not already regulated, 4 were identified as potential quarantine pests for regulation (Table 3), of those, 2 were added to the CFIA’s regulated pest list and one was designated for additional review.

**Figure 1.**
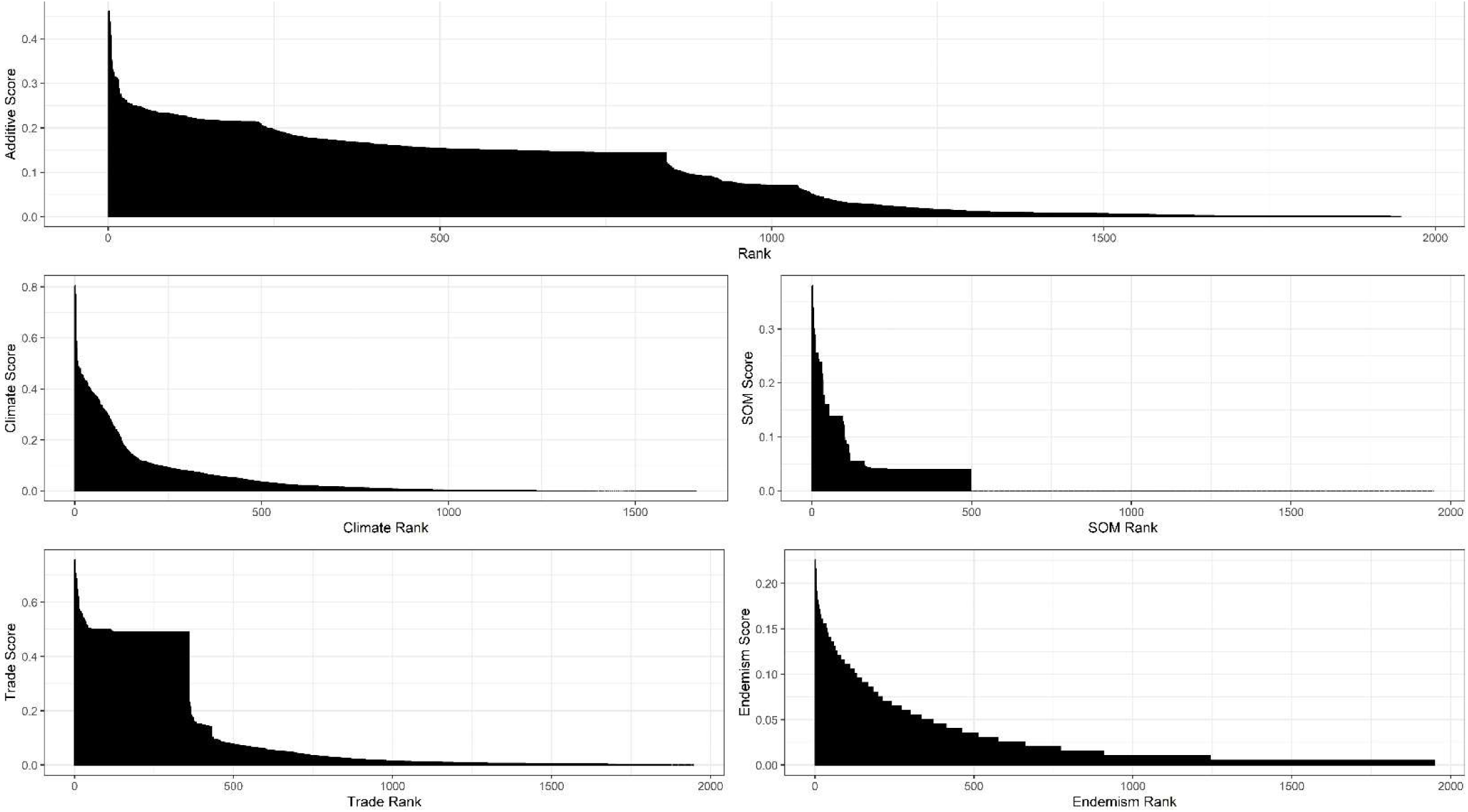
Score-rank relationships for total score, and its components. Includes species identified in either Climatch or SOM results.

**Table 2.** Wood-boring beetle short-list identified by horizon scanning.

| Rank | Family | Species | # Records | # Countries | Additive Score | Categorization Result | Regulatory Result |
| --- | --- | --- | --- | --- | --- | --- | --- |
| 1* | Cerambycidae | *Monochamus sutor* | 6419 | 32 | 0.462 | Already regulated |  |
| 2 | Cerambycidae | *Trichoferus campestris*<br>(*Hesperophanes campestris*) | 310 | 22 | 0.447 | Already regulated |  |
| 3 | Cerambycidae | *Monochamus scutellatus* | 7481 | 2 | 0.438 | Already regulated |  |
| 4 | Cerambycidae | *Monochamus notatus* | 1657 | 2 | 0.408 | Already regulated |  |
| 5 | Curculionidae | *Hylesinus varius* | 8792 | 27 | 0.353 | Potential quarantine pest | Added to regulated pest list |
| 6 | Cerambycidae | *Monochamus marmorator* | 137 | 2 | 0.349 | Already regulated |  |
| 7* | Cerambycidae | *Monochamus urussovii* | 281 | 10 | 0.330 | Already regulated |  |
| 8* | Cerambycidae | *Anoplophora glabripennis* | 129 | 12 | 0.324 | Already regulated |  |
| 9 | Curculionidae | *Trypodendron signatum* | 4151 | 17 | 0.323 | Not potential quarantine pest |  |
| 10 | Cerambycidae | *Callidium aeneum* | 1699 | 20 | 0.315 | Not potential quarantine pest |  |
| 11 | Curculionidae | *Ips typographus* | 12715 | 24 | 0.314 | Already regulated |  |
| 12 | Buprestidae | *Trachys minutus* | 5979 | 33 | 0.313 | Not potential quarantine pest |  |
| 13 | Curculionidae | *Pityogenes bidentatus* | 4186 | 19 | 0.312 | Not potential quarantine pest |  |
| 14 | Cerambycidae | *Monochamus clamator* | 736 | 4 | 0.311 | Already regulated |  |
| 15* | Cerambycidae | *Monochamus galloprovincialis* | 3745 | 31 | 0.309 | Already regulated |  |
| 16 | Cerambycidae | *Monochamus titillator* | 82 | 2 | 0.288 | Already regulated |  |
| 17 | Buprestidae | *Agrilus planipennis* | 2095 | 5 | 0.285 | Already regulated |  |
| 18 | Cerambycidae | *Monochamus carolinensis* | 659 | 2 | 0.275 | Already regulated |  |
| 19 | Cerambycidae | *Chlorophorus annularis* | 245 | 31 | 0.275 | Not potential quarantine pest |  |
| 20 | Curculionidae | *Ips sexdentatus* | 3785 | 18 | 0.268 | Potential quarantine pest | Added to regulated pest list |
| 21 | Brentidae | *Cylas formicarius* | 344 | 36 | 0.268 | Potential quarantine pest | Not added to regulated pest list |
| 22 | Cerambycidae | *Agapanthia villosoviridescens* | 21711 | 36 | 0.265 | Not potential quarantine pest |  |
| 23 | Curculionidae | *Cnestus mutilatus* | 18 | 1 | 0.265 | Potential quarantine pest | Under review |
| 24 | Cerambycidae | *Anoplophora chinensis* | 585 | 12 | 0.263 | Already regulated |  |

\*Regulated at genus level in Canada.

**Table 3.**
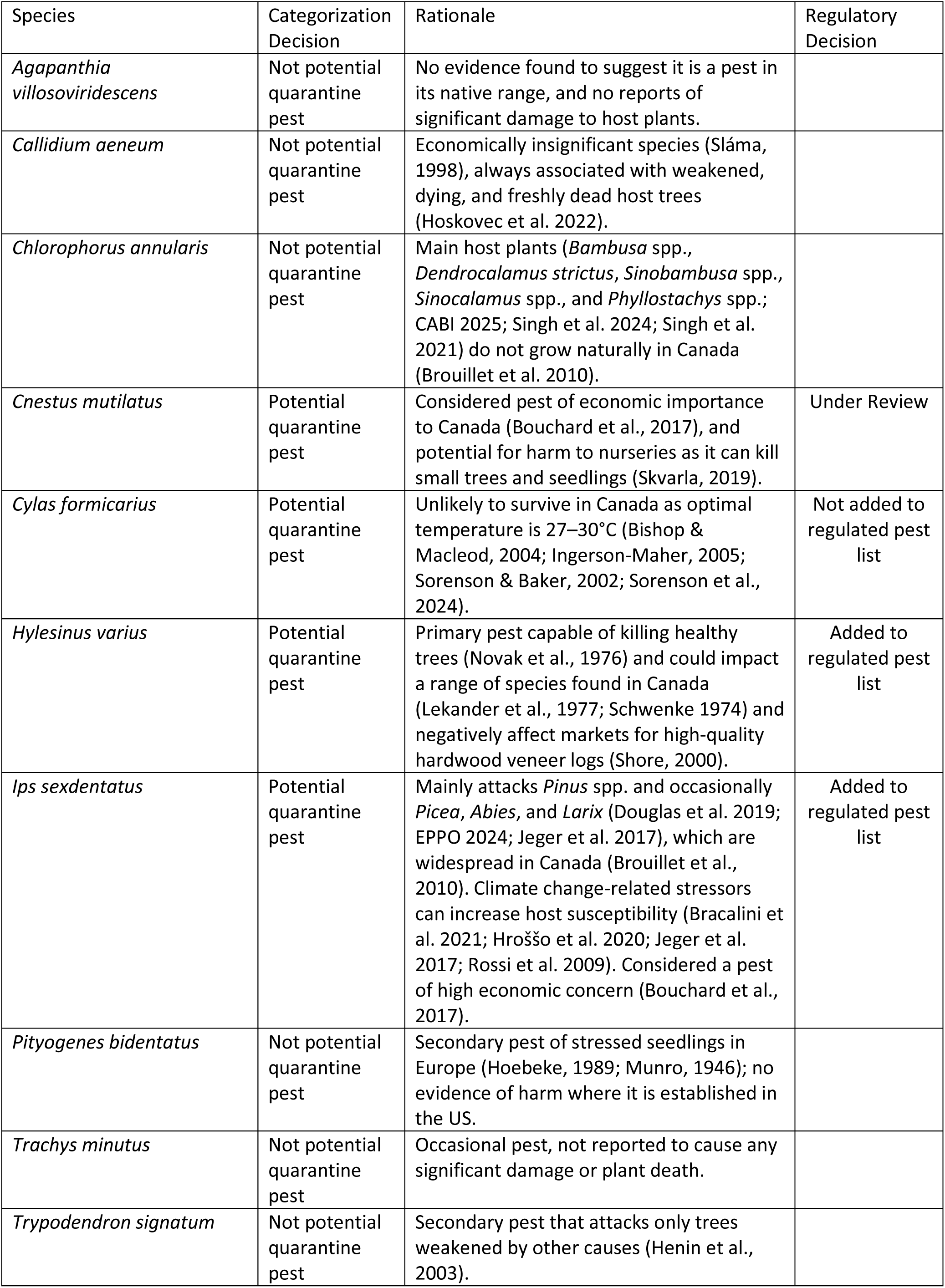
Short-listed species, categorization decisions, and rationale.

Overlaying climate suitability maps can identify potential hotspots for pest establishment, which shows hotspots of climate suitability for wood-boring beetles in areas of western, southern, and eastern Canada (Figure 2).

**Figure 2.**
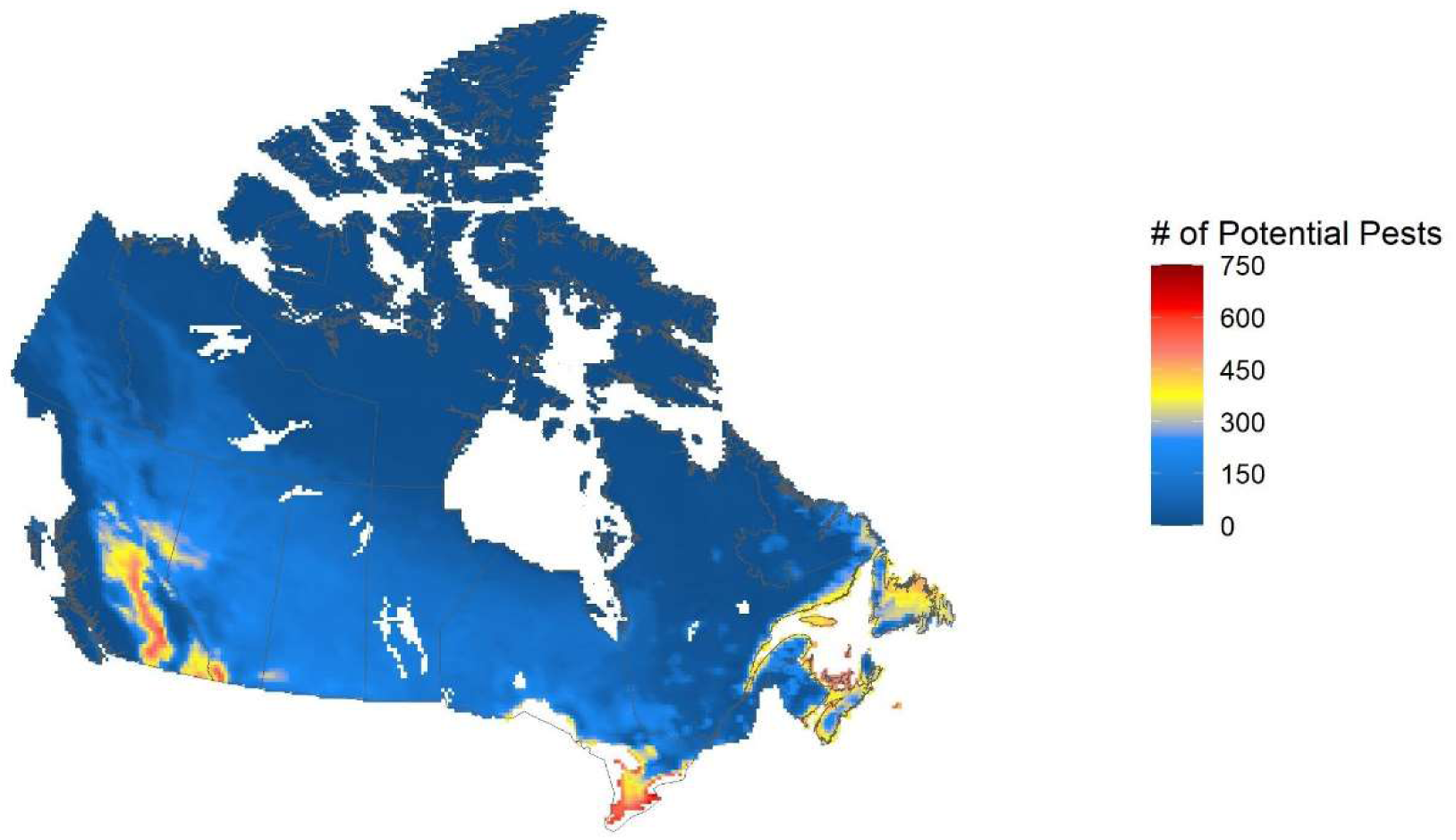
Number of species from the horizon scan with climate suitability in Canada. Redder areas indicate a greater number of species having climate suitability.

Looking at scores across families, Bostrichidae, Buprestidae, and Curculionidae species tended to have higher scores than other families, while Anobiidae and Brentidae species tended to have lower scores (Figure 3).

**Figure 3.**
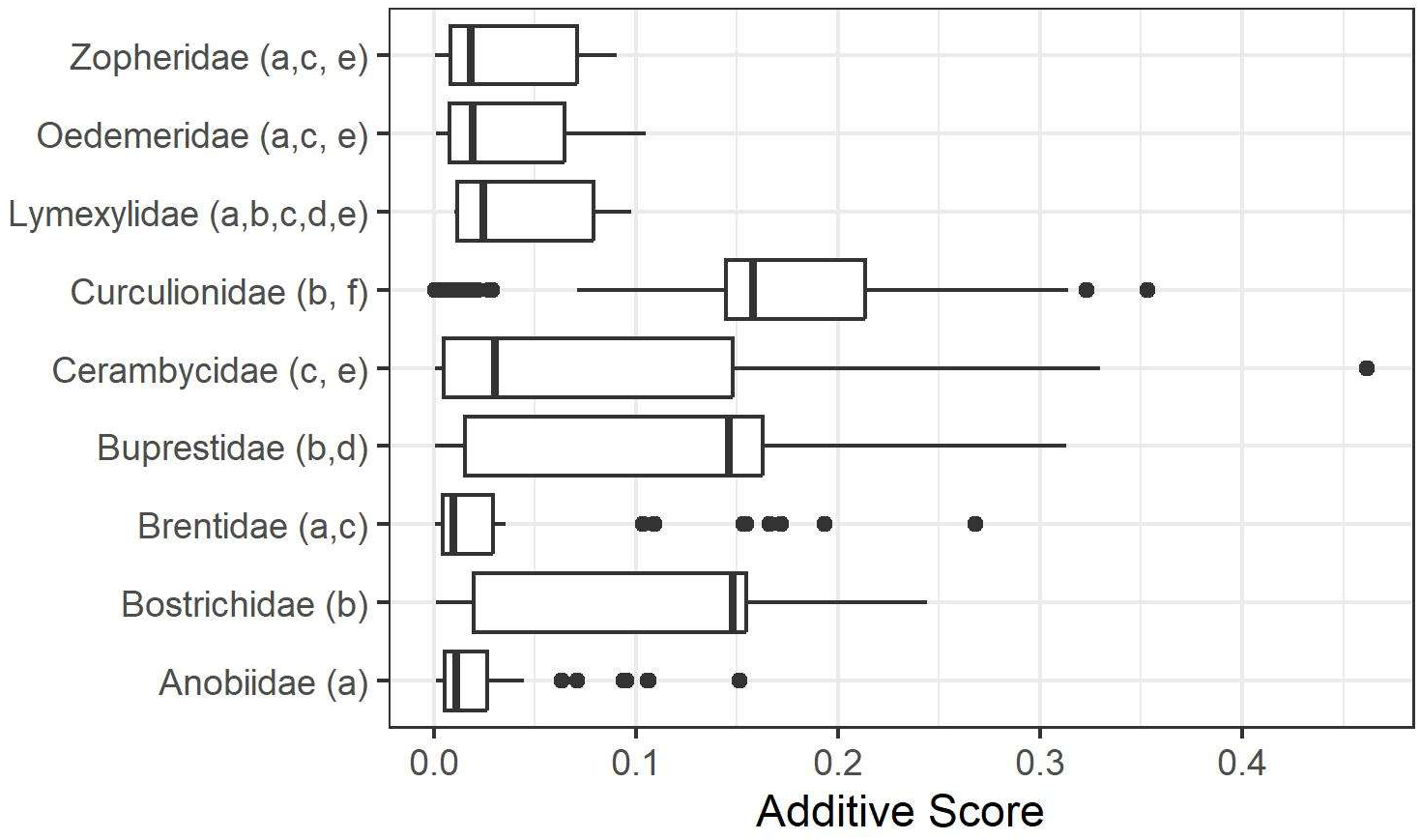
Scores of species vary by family. Identical letters indicate scores are statistically equivalent.

## Discussion

Horizon scanning is a necessary (Antoniou et al., 2024) but time-consuming part of tasks conducted by NPPO risk assessors. Automation of parts of it can alleviate repetitive aspects of this task, while removing some subjectivity inherent in decision-making. Quantitative horizon scanning with climate matching, machine learning, and risk assessment criteria, produced lists of potential pests for further regulatory evaluation that are consistent with existing procedures, and led to species being added to Canada’s regulated pest list. This suggests that the methods used here are valuable for proactive identification of potential pests for regulation by NPPOs and may support Target 6 of the Kunming-Montreal Global Biodiversity Framework to reduce the introduction of IAS (UN, CBD 2022) via proactive regulation.

*Cnestus mutilatus, Cylas formicarius, Hylesinus varius* and *Ips sexdentatus* were identified as potential quarantine pests, and both *Hylesinus varius* and *Ips sexdentatus* were added to Canada’s regulated pest list because of our analysis. While the relatively low number of species that were identified as meeting the biological criteria to be considered potential quarantine pests in follow-up categorizations might suggest the outcome is disappointing, it would be disconcerting if the opposite were true: if all species identified were categorized as potential quarantine pests, or very many, that could indicate rank cut-offs were too high. As such, 75% of the short-listed species were either already regulated in a Canadian jurisdiction or were considered a potential quarantine pest. The relatively high proportion of non-regulated species flagged by this method that were determined not to be potential quarantine pests (60%) points to the value of literature-based evaluations once quantitative horizon scanning has narrowed the scope of the problem in a resource-constrained regulatory context.

Previous horizon scans identified lists of 40 or more candidate species (e.g. Lieurance et al., 2023), in some cases exceeding 100 (e.g. Kenis et al., 2022). From a regulatory point of view, such lists can be impracticably long, since each species must undergo a categorization and possibly full risk assessment, as well as pest risk management. Our methods here reduce this issue, using a range of scoring criteria that are already used in formal risk assessments, generating risk pre-assessments to guide responses to potential pests. Through this process, we generated a short-list of 10 species for further consideration (excluding already-regulated species), an approximately 4-fold reduction in candidate species compared to previous methods and have identified 4 species that were potential quarantine pests for Canada out of a pool of thousands. Given that IAS are expected to continue to accumulate and accelerate with climate change (Seebens et al., 2021), prospective methods can help support regulations intended to reduce the likelihood that new pests are introduced. Early identification of pests allows for more rapid implementation of management activities that can limit entry and improve surveillance. Our methods could be applied to regulated pests to identify higher priority species for surveys and inspections, and to support decisions related to optimizing surveillance resources among pests.

Results can be compared to interception and establishment records in the literature. Interception frequency of true bark beetles (Scolytinae) is highly correlated with establishment frequency (Brockerhoff et al., 2006). Members of the Scolytinae, Cerambycidae, and Curculionidae (non-Scolytinae) figured prominently in an evaluation of wood-boring beetle interceptions by the US (Haack, 2006). 20 of the 24 short-listed species are from highly-intercepted families, suggesting that not only are these species higher risk, but they are also more likely to be present on imported products. While our analysis suggests Curculionidae are associated with higher risk than other families, 14 of 24 short-listed species were from the Cerambycidae. This suggests our approach is valuable for identifying potential pests, but not necessarily for identifying general patterns across taxa, possibly due to country-specificity of the input data.

## Drawbacks and limitations

Programmatically accessible data sources are a primary limitation on quantitative horizon scanning. Regarding trade data, the data were at the country-level, which poses some caveats with respect to geographic resolution of trade data but allows for broader applicability to other jurisdictions that may only track trade at a country-level.

The nine families used to identify wood-boring beetle species were selected based on characters for each entire family, therefore any member of a family may be identified, not just known wood borers. That is, the final list may contain beetles that are not wood borers.

The SOM was run on the taxa of interest. Due to dependency of wood-boring beetles on suitable hosts, it may be valuable to add hosts to the SOM. This could help in cases where a target species is known to occur in only one administrative region, which the current method is unable to resolve since such species are not shared between regions.

## Conclusions

Quantitative horizon scanning is a suitable method for identifying potential invasive species that may require regulation to prevent their invasion.

## Acknowledgements

We would like to thank Dominique Pelletier, Arvind Vasudevan, Angela Murray and Josh Persi for reviewing the manuscript.

## Supplementary Information

### Climate Data

**Table S1.** Climate variables considered for climate matching.

| CHELSA Name | Description | Included in Climatch Analysis? |
| --- | --- | --- |
| Bio1 | mean annual air temperature |  |
| Bio5 | mean daily maximum air temperature of the warmest month | Yes |
| Bio6 | mean daily minimum air temperature of the coldest month | Yes |
| Bio8 | mean daily mean air temperatures of the wettest quarter |  |
| Bio9 | mean daily mean air temperatures of the driest quarter |  |
| Bio10 | mean daily mean air temperatures of the warmest quarter |  |
| Bio11 | mean daily mean air temperatures of the coldest quarter |  |
| Bio12 | annual precipitation amount |  |
| Bio13 | precipitation amount of the wettest month |  |
| Bio14 | precipitation amount of the driest month |  |
| Bio16 | mean monthly precipitation amount of the wettest quarter | Yes |
| Bio17 | mean monthly precipitation amount of the driest quarter | Yes |
| Bio18 | mean monthly precipitation amount of the warmest quarter | Yes |
| Bio19 | mean monthly precipitation amount of the coldest quarter | Yes |

**Figure S1.**
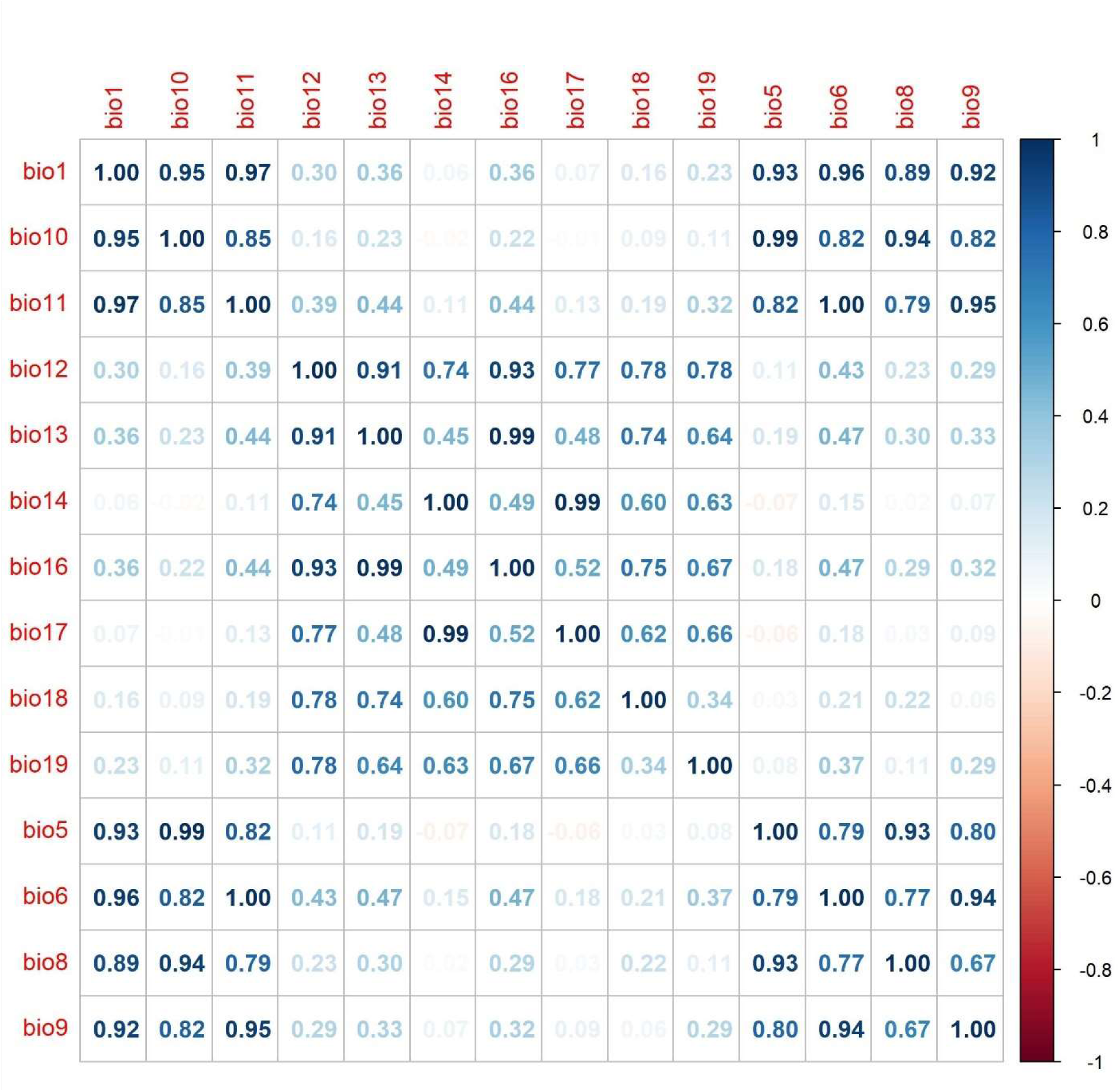
Full correlation plot for all bioclimatic variables.

**Figure S2.**
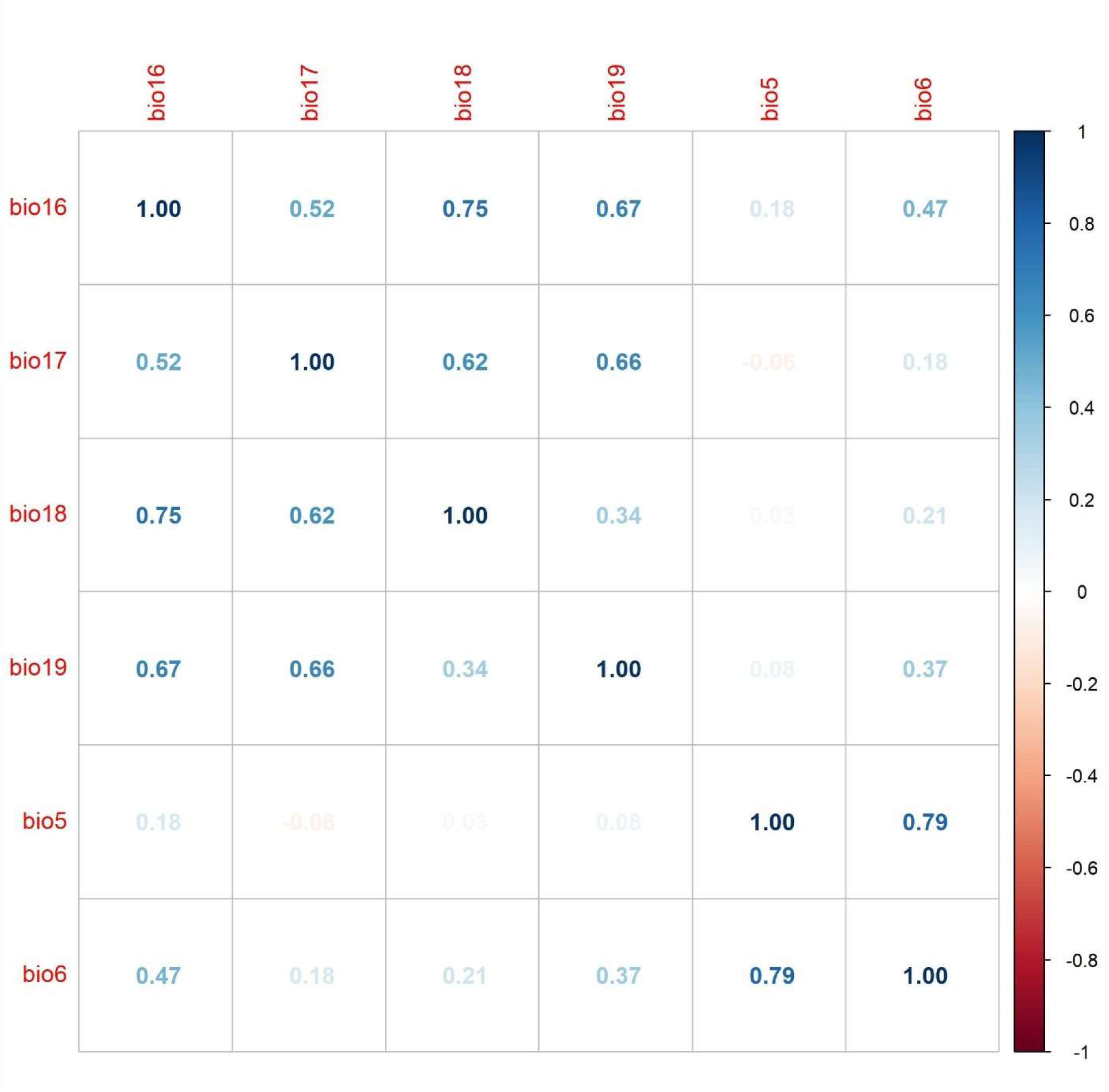
Final correlation plot for the bioclimatic variables used in the present study.

